# GFP-Forked, a genetic reporter for studying *Drosophila* oocyte polarity

**DOI:** 10.1101/449090

**Authors:** Raju Baskar, Anna Bakrhat, Uri Abdu

**Affiliations:** Department of Life Sciences, Ben-Gurion University of the Negev, Be’er Sheva, Israel.

**Keywords:** *Drosophila*, Forked, Oocyte polarity, Microtubules, ncMTOC

## Abstract

The polarized organization of the *Drosophila* oocyte can be visualized by examining the asymmetric localization of mRNAs, which is supported by networks of polarized microtubules (MTs). In this study, we used the gene *forked*, the putative *Drosophila* homologue of *espin*, to develop a unique genetic reporter for asymmetric oocyte organization. We generated a null allele of the *forked* gene using the CRISPRCas9 system and found that *forked* is not required for determining the axes of the *Drosophila* embryo. However, ectopic expression of a truncated form of GFP-Forked generated a distinct network of asymmetric Forked, which first accumulated at the oocyte posterior and was then restricted to the anterolateral region of the oocyte cortex in mid-oogenesis. This localization pattern resembled that reported for the polarized MTs network. Indeed, pharmacological and genetic manipulation of the polarized organization of the oocyte showed that the filamentous Forked network diffused throughout the entire cortical surface of the oocyte, as would be expected upon perturbation of oocyte polarization. Finally, we demonstrated that Forked associated with Short-stop and Patronin foci, which assemble non-centrosomal microtubule-organizing centers. Our results thus show that clear visualization of asymmetric GFP-Forked network localization can be used as a novel tool for studying oocyte polarity.

**Summary statement:** The novel asymmetric Forked network could be used as a genetic reporter for visualizing and studying oocyte polarity.

## Introduction

Correct localization of intracellular messenger RNAs (mRNAs) encoding morphogenetic proteins to their distinct subcellular domains are crucial for specification of the body axes of the *Drosophila* embryo. The roles of three major asymmetrically localized mRNAs, *gurken (grk), bicoid (bcd),* and *oskar (osk)*, in the process of establishing axial patterning of the oocyte and embryo in mid-oogenesis were clearly established. *grk* is localized around the oocyte nucleus and determines the dorsal-ventral axis of the oocyte and embryo (Gonzalez-Reyes et al., 1995; Neuman-Silberberg and Schupbach, 1993; Neuman-Silberberg and Schupbach, 1996; Roth et al., 1995), whereas *bcd* is localized to the extreme anterior of the oocyte and determines the anterior pattern of the embryo upon translation (Driever and Nusslein-Volhard, 1988a; Driever and Nusslein-Volhard, 1988b). At the same time, *osk* is localized to the posterior of the oocyte and initiates the development of future germ cells and the embryo abdomen (Ephrussi et al., 1991).

Asymmetric mRNA localization during mid-oogenesis depends on microtubules (MTs), actin networks and motor proteins. During mid-oogenesis (i.e., stages 9 and 10), MTs within the oocyte are organized asymmetrically, with non-centrosomal microtubule-organization centers (ncMTOCs) being localized solely to the anterior and lateral cortexes of the oocyte (Huynh and St Johnston, 2004; Nashchekin et al., 2016; Theurkauf, 1994). Regulation of the asymmetrical oocyte MT network is mainly controlled by two sequential processes. Initially, an MTOC positioned at the posterior end of the oocyte, close to the nucleus, which is in a symmetric position at this stage, is established. Signaling from *grk* to EGF receptors at the posterior end of the oocyte (stage 6-7) establishes posterior follicle cell fate (Gonzalez-Reyes et al., 1995; Roth et al., 1995). Subsequently, an unknown signal produced by posterior follicle cells leads to the establishment of Par-1 kinase activity in the posterior oocyte cortex, thereby defining the antero-lateral versus posterior cortical domain in the oocyte (Doerflinger et al., 2006; Shulman et al., 2000). In addition, the same unknown signal triggers disassembly of the posterior MTOC. At the same time, nucleation of new MTs at the oocyte anterior end results in a reversal of polarity within the oocyte (Gonzalez-Reyes et al., 1995; Roth et al., 1995; Theurkauf et al., 1992). Par-1 kinase now restricts the polarization of MTs to the antero-lateral region of the oocyte by suppressing MT nucleation at the oocyte posterior end (Doerflinger et al., 2006; Parton et al., 2011).

Visualization of the asymmetric organization of the *Drosophila* oocyte is achieved either by *in situ* hybridization or by staining with antibodies directed against several localized mRNA, such as *grk*, *bcd* and *osk*. In addition, the polarized organization of oocyte MTs can be demonstrated either by staining using anti-tubulin antibodies or by the expression of MT-associated proteins fused to GFP. Detection of the polarized MT network using anti-tubulin antibodies, however, requires the use of a special protocol (Legent et al., 2015).

In studying the role of *forked* gene, the *Drosophila* homologue of *espin*, in oogenesis, we discovered that ectopic expression of GFP-Forked protein could be used as a novel tool for analyzing oocyte polarity. First, we demonstrated that oocytes containing mutations in *forked* showed no defects in polarity. On the other hand, upon over-expression of GFP-Forked, the protein first accumulated at the oocyte posterior end and was then restricted to the anterolateral region of the oocyte cortex in mid-oogenesis. We showed that this unique asymmetric localization of GFP-Forked depends on the polarized microtubule network. We further found that ectopic expression of Forked is associated with Short-stop and Patronin foci. Thus, our results reveal that the novel asymmetric Forked network can be used as a genetic reporter for visualizing and studying oocyte polarity.

## Results

### Forked is not required for oocyte axis determination

Previous work from our lab and others revealed similarities between oocyte and bristle cytoskeleton organization (Abdu et al., 2006; Amsalem et al., 2013; Bitan et al., 2010; Dubin-Bar et al., 2011; Dubin-Bar et al., 2008; Lundh et al., 2011; Otani et al., 2015; Shapiro and Anderson, 2006). It was shown that the complex of proteins containing Spn-F, Ik2 and Jvl plays a similar role in both oocyte and bristle actin network organization. Generating actin bristle bundles requires two actin-bundling proteins, Singed (sn), the *Drosophila* Fascin homologue, and Forked, the putative *Drosophila* homologue of *espin*. Previously, it was shown that Singed is required for the formation of cytoplasmic actin bundles in nurse cells (Cant et al., 1994). However, the role of *forked* in oogenesis is still unknown.

Since the *f*^*36a*^ (Hoover et al., 1993) allele failed to eliminate all *forked* splice forms and there is no molecular characterization for the *f*^*15*^ allele containing a stop codon at Q206 (flybase; FBrf0191634), we decided to delete the *forked* gene using CRISPR/Cas9 technology. For this, we generated the null allele so as to eliminate all alternatively splice forms of *forked* (Fig. 1E). Indeed, our PCR analysis revealed that we replaced the entire *forked* genomic region with dsRed (Fig. 1F). Moreover, homozygous alleles showed aberrant *forked* bristle phenotype (Fig. 1H). Moreover, we found that *forked* null allele females were fertile (Table S1).

**Figure 1:**
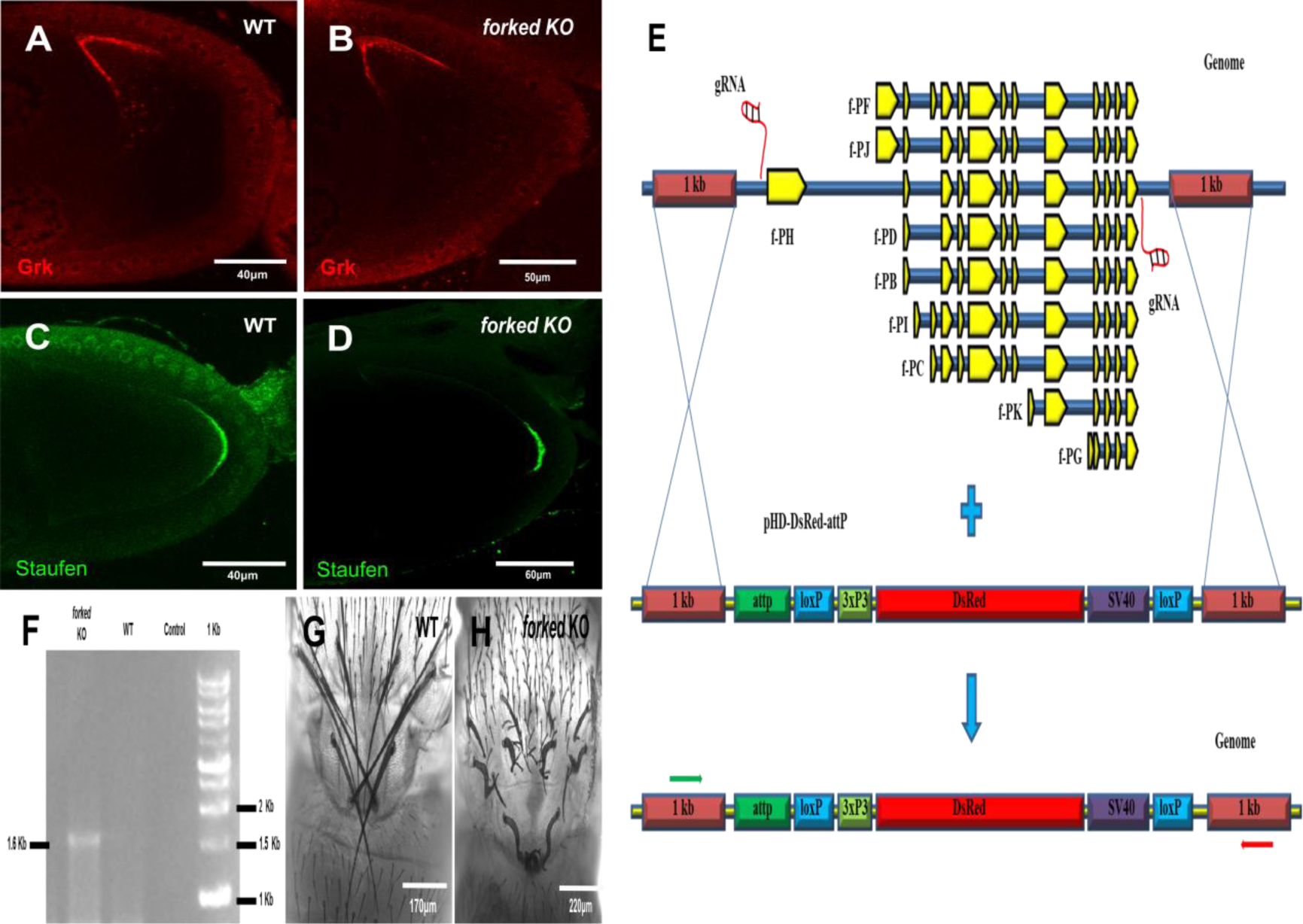
Forked is not required for oocyte axis determination. Confocal images of stage 9 egg chamber from flies of WT (A, C) and CRISPR *forked KO* (B, D) stained with Grk antibody in red (A, B) and stained with Staufen in green (C, D). (E) Schematic diagram showing the nature of *forked* locus and generation of *forked KO* (null allele) using CRISPR mediated HDR with the donor vector of pHDDsRed-attP. All the nine *forked* isoforms exons are shown in yellow on locus which is in blue color. Green and Red arrows represent the binding sites of forward and reverse primer that were used for genotyping. (F) PCR analysis of genotyping of *forked* knock out (KO). The size of 1.6 Kb band in *forked* KO lane depicts the complete deletion of *forked* from the genome and replaced with DsRed using the primers are shown in (E) green and red arrows. Confocal images of thorax region from the pharate adult of WT (G) and *forked* KO (H).

Next, we tested whether *forked* plays a role in oocyte polarity. As a read-out for any defect in oocyte polarity, we tested the localization of Gurken (Grk), which is responsible for determination of the dorsal-ventral axis of the egg (Gonzalez-Reyes et al., 1995), and Staufen (Stau), which is required for the localization of both the anterior axis determinants Bicoid (Bcd) and the posterior axis determinant Oskar (Osk) (St Johnston et al., 1991). We found that *forked*-deleted flies presented no obvious defects in terms of the localization of either Grk or Stau (Fig. 1B, D; n=20, 100%).

### Forked marks a distinct asymmetric network in the oocyte

Although our results showed that Forked is not required for oocyte polarity, we tested the localization pattern of ectopic GFP-Forked expression. Since it had been shown that extra copies of the *forked* gene affected bristle development (Petersen et al., 1994), we generated a truncated form of the *forked* gene fused to DNA encoding GFP (see Material and Methods). We found that although the GFP-Forked chimera was localized to bristle actin bundles (Fig. S1A-C), it failed to rescue the *forked* bristle phenotype (Fig. S1E-F). Next, when we analyzed the localization of GFP-Forked in the egg-chamber. To our surprise, we found that the ectopic expression of GFP-Forked presented a unique localization pattern. At stage 5, GFP tagged-Forked decorated a filamentous network that radiated from the circumferential cortical posterior surface of the oocyte towards the oocyte cytoplasm (Fig. 2C, D). However, in stage 9 egg chambers, this localization pattern had changed, such that the GFP tagged-Forked filamentous network was now restricted to the anterolateral cortex of the oocyte (Fig. 2E-G) and was completely absent from the posterior end of the oocyte (Fig. 2H). Moreover, at stage 10, this asymmetrical localization pattern of GFP-Forked could still be detected (Fig. 2I-L). We tested for effects of GFP-Forked expression and found that over-expression of GFP-Forked had no effect on female fertility (Table S2). We then asked whether this asymmetric filamentous network co-localized with actin. Using the general actin marker phalloidin, we found that the filamentous asymmetric network decorated by Forked indeed co-localized with actin (Fig. 3A-F). To summarize, our results suggest that ectopic expression of Forked generates a distinct filamentous network in the oocyte.

**Figure 2:**
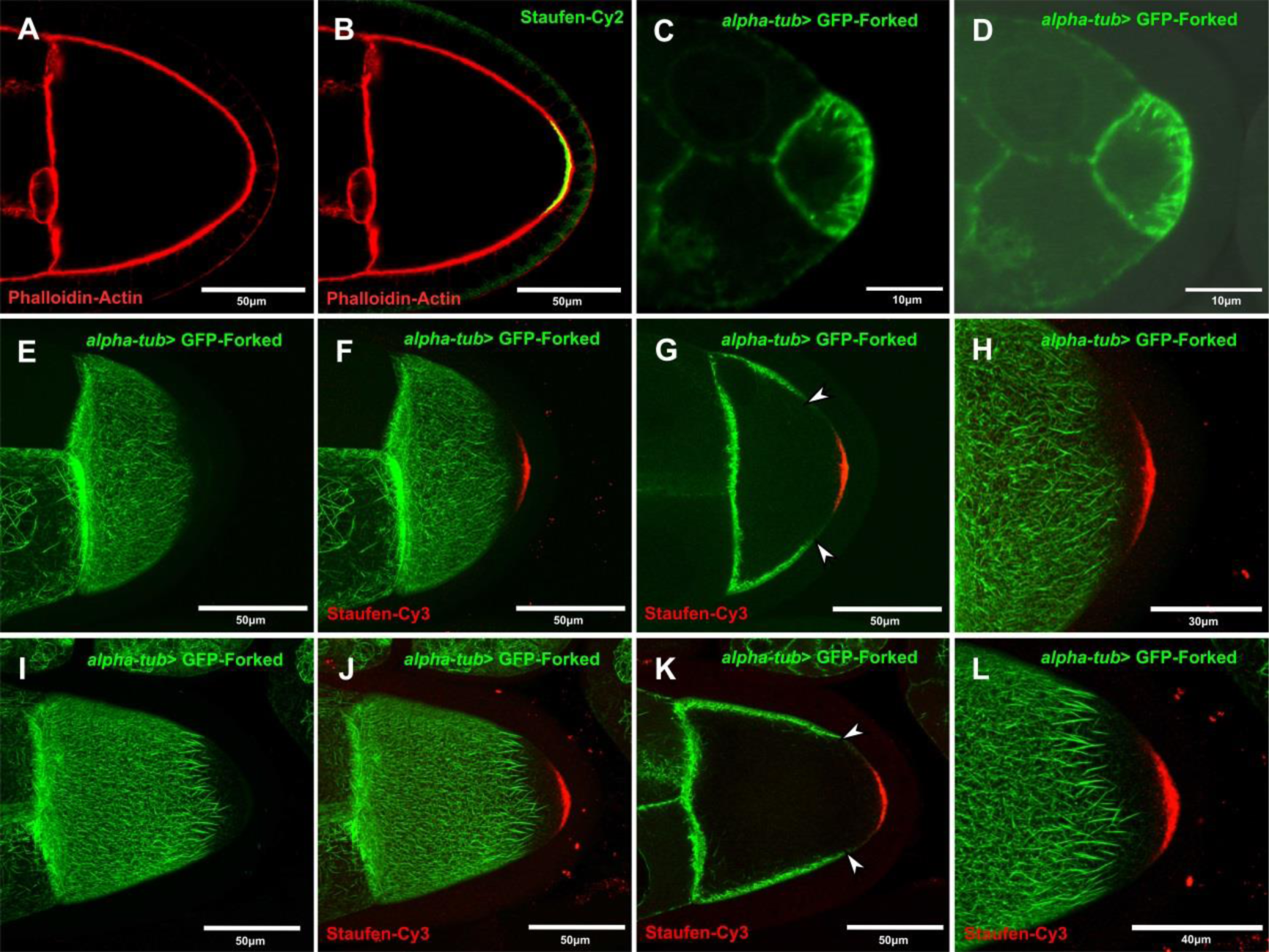
Forked marks a distinct asymmetric network in oocyte. Confocal images from (A, B) Wild type stage 9 egg chamber stained with phalloidin in red and staufen antibody in green (B), (C-D) *alpha-tub>* GFP-Forked stage 5, stage 9 (E-H) and stage 10 (I-L) egg chambers stained with staufen-Cy3 antibody. Images (E, F, H) and (I, J, L) are Confocal Z-series projections. (G) and (K), are single confocal slice from the Z-series projections of image (F) and (J), respectively. D is merge image of C with DIC. Arrows in (G) and (K) are pointing towards the limit of the asymmetric network marked by GFP-Forked.

**Figure 3:**
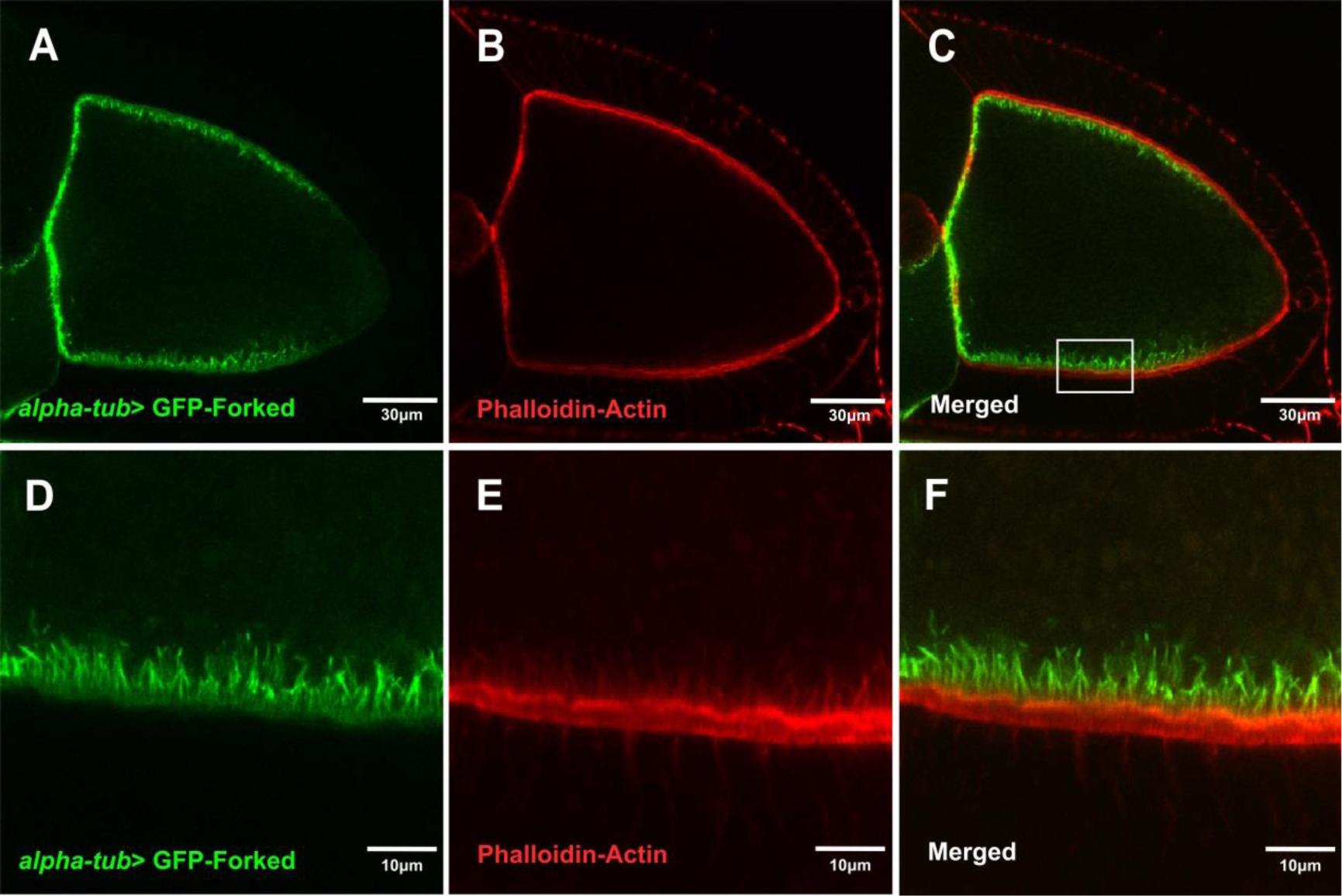
Co-localization of Forked-asymmetric network with actin marker. (A-C) Confocal image of stage 10 egg chamber expressing *alpha-tub>* GFP-Forked, stained with phalloidin for actin in red. (D-F) Enlargement of boxed region of antero-lateral segment from (C).

### The asymmetric Forked network depends on microtubules

We asked whether this decorated filamentous and asymmetric Forked network depended on MTs. For this purpose, flies expressing GFP-tagged Forked in the germline were fed with the microtubule-depolymerizing agent colchicine. We found that feeding flies with colchicine affected MT organization, as reflected by the mis-localization of the oocyte nucleus (Fig. 4B, D). Moreover, the decorated filamentous Forked network was no longer organized in an asymmetrical manner in the egg chambers, instead being diffusely distributed throughout the oocyte cortical surface (n=40, 100%; Fig. 4A, C,). This indicates that the filamentous asymmetric Forked network is dependent on MT organization.

**Figure 4:**
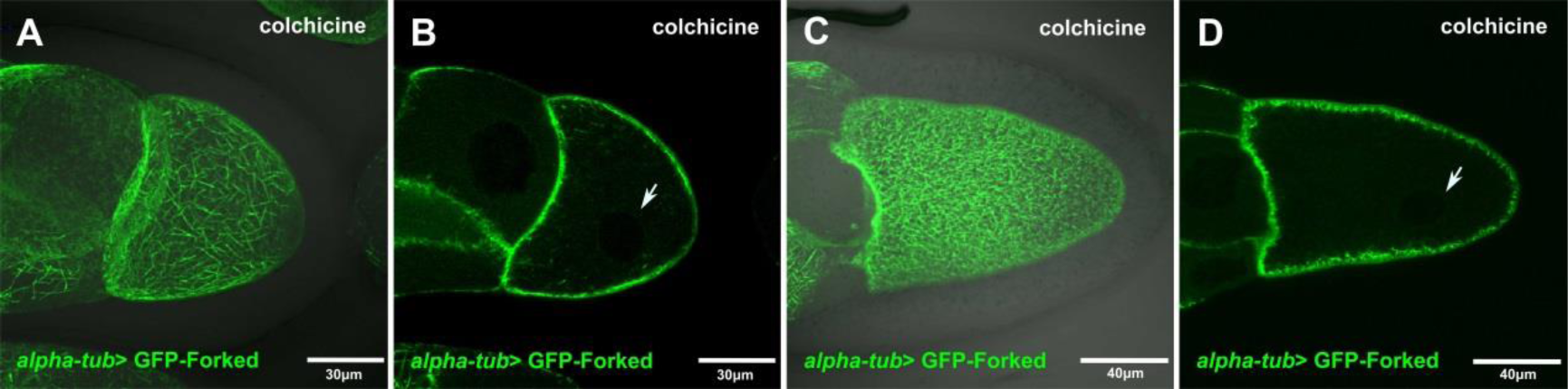
The asymmetric Forked network depends on microtubules. (A, B) Confocal images from stage 9 and stage 10 (C, D) egg chambers expressing *alpha-tub>* GFP-Forked flies, fed with colchicine. Where (A) and (B) are Confocal Z-series projections. (B) and (D), are single confocal slice from the Z-series projections of image (A) and (C) which was also merged with DIC, respectively. Arrows in (B) and (D) are marking the oocyte nucleus.

### The asymmetric Forked network depends on a polarized microtubule network

Next, we tested whether the novel asymmetrical Forked network requires a polarized microtubule network. We, therefore, first analyzed the localization pattern of GFP-tagged Forked in a *grk* mutant background, where during mid-oogenesis the MTs fail to repolarize and the oocyte nucleus often fails to migrate. We found that in stage 9 egg chambers from *grk* mutants (n=45, 100%), the Forked filamentous network was diffusely localized throughout the cortical surface of the entire oocyte (Fig. 5A) and was no longer restricted to the anterolateral cortex, as in the wild type (WT) (Fig. 5B). Moreover, at stage 10 (n=20, 25%; Fig. 5C), the filamentous Forked network generated a cage-like structure around the mis-localized oocyte nucleus (Fig. 5C).

**Figure 5:**
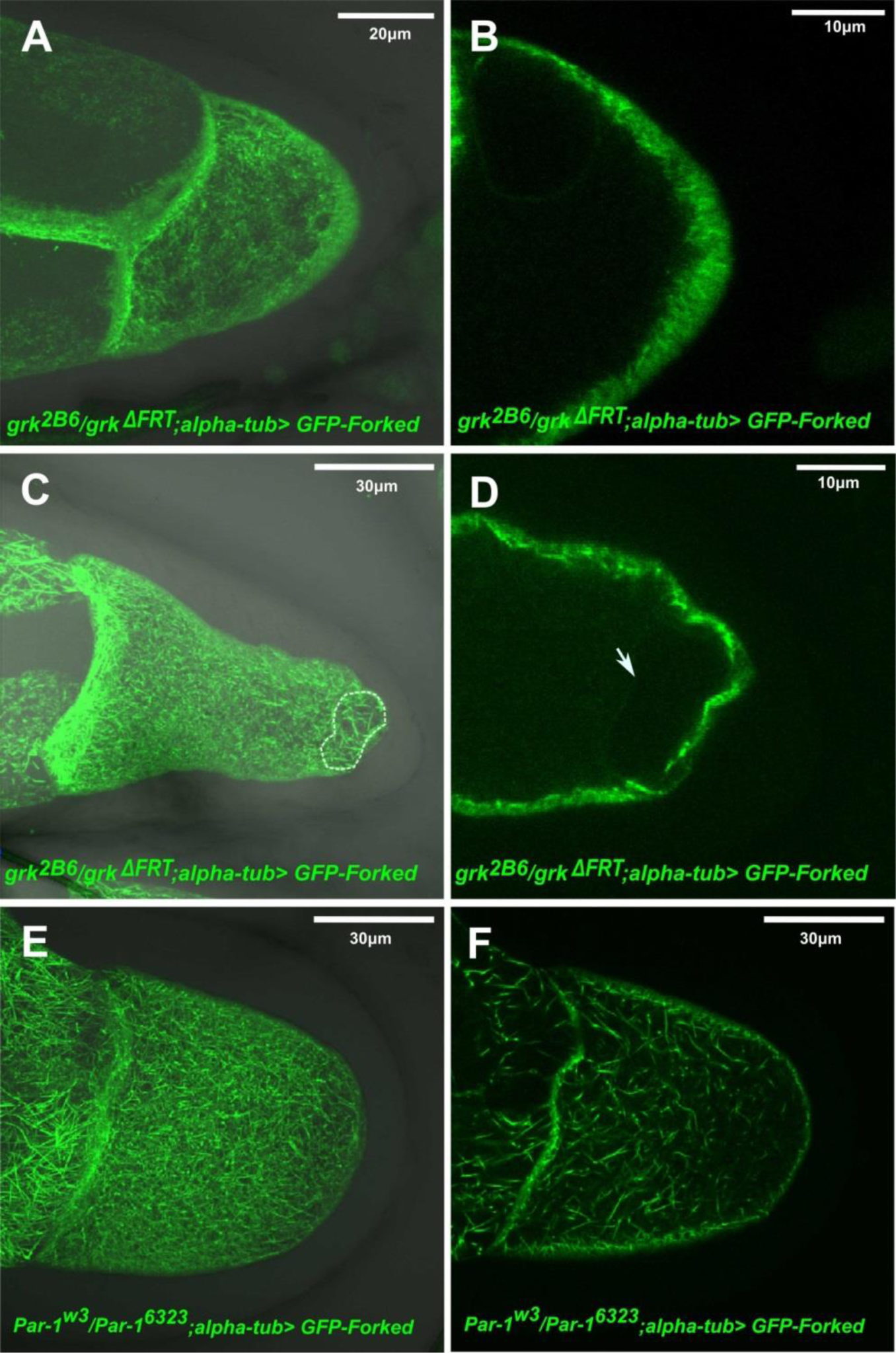
The asymmetric Forked network depends on polarized microtubule network. Confocal Z-series projections of stage 9 (A, B) and stage 10 (C, D) egg chamber from *grk* mutant flies expressing *alpha-tub>* GFP-Forked, merged with DIC. (B) and (D) are single confocal slice from the Z-series projections of image (A) and (B), respectively. White dot lines represent the cage around the nucleus (C) and arrow in (D) points to the oocyte nucleus. (E, F) Confocal Z-series projections merged with DIC of stage 10 (E) egg chamber from *Par-1* mutants flies expressing *alpha-tub>* GFP-Forked. (F) Single confocal slice from the Z-series projections of image E.

Since it has been reported that PAR-1 is required for polarized MT organization in the oocyte by preventing MT nucleation sites from forming at the posterior cortex (Parton et al., 2011), we analyzed the localization pattern of GFP-tagged Forked in a *par1* hypomorph mutant background that allows egg chambers to progress to mid-oogenesis (Parton et al., 2011; Shulman et al., 2000). In stage 9-10 egg chambers of *par1* mutant females, the filamentous Forked network was no longer organized in an asymmetrical manner, instead being diffusely distributed throughout the entire cortical surface of the oocyte (Fig. 5E, F, n=35, 100%).

Another factor required for organizing the polarized MT network in the oocyte is the actin-microtubule cross linker *short stop* (*shot*). Recently, it was shown that *shot*, along with *patronin*, a *Drosophila* microtubule minus-end-binding protein, which encodes a CAMSAP homologue, are both required for MT organization in the oocyte by assembling non-centrosomal MTOCs at the antero-lateral cortex of the oocyte (Nashchekin et al., 2016). Since, *shot* mutant egg chambers fail to develop past earlier stage of oogenesis, we used RNAi expressed in the germline via the UAS GAL4 system to down-regulate levels of the *shot* transcript. We found in *shot* knockdown egg chambers, the filamentous Forked asymmetric network was also diffusely distributed throughout the cortical surface of the oocyte (n=20, 100%; Fig. S2C, D). In summary, we observed that whenever the polarized microtubule network was disrupted, the filamentous Forked network also lost polarity. Therefore, the asymmetric Forked network depends on a polarized microtubule network.

### Forked is associated with ncMTOCs

Recently, it was shown that the ncMTOCs, which comprises Shot and Patronin, localizes to the anterolateral region of the oocyte (Nashchekin et al., 2016). Since the asymmetric localization pattern of Forked resembles that of Shot and Patronin, we assessed whether the ectopic expression of Forked generated this asymmetric network with ncMTOCs. First, we checked whether this asymmetric network co-localized with Shot. Using the UAS-Gal4 system, we expressed mCherry-Forked in the background of flies endogenously expressing Shot-YFP. We found that Shot-YFP co-localized with the Forked-asymmetric network (Fig. 6 A-C′′′). Closer examination revealed that Forked was associated with Shot foci along the anterolateral cortex. Then, we considered whether disruption of the polarized microtubule network with colchicine was responsible for the change in Forked localization. Accordingly, flies expressing both mCherry-Patronin and GFP-Forked in the germline were fed with colchicine. We found that similar to the diffuse pattern of Forked (Fig. 6E-F) seen upon such drug treatment, Patronin foci were diffusely distributed throughout the oocyte, when compared to untreated animals (n=30, 100%; Fig. 6D), and co-localized with Forked (Fig. 6G-I). Since we found that the asymmetric Forked network associated with ncMTOCs, we asked whether ncMTOCs is also associated with actin-enriched aggregates in mutants with abnormal actin networks, such as *ik2, spn-F* and *jvl* mutants (Abdu et al., 2006; Dubin-Bar et al., 2011; Shapiro and Anderson, 2006),. We found that mCherry-Patronin was associated with ectopic actin structures in both *ik2*-RNAi (Fig. 6J-O) and *jvl* mutants (data not shown). Thus, these results suggest that the ectopic expression of Forked in the ovary generates an asymmetric network, which emanates from ncMTOCs in the oocyte.

**Figure 6:**
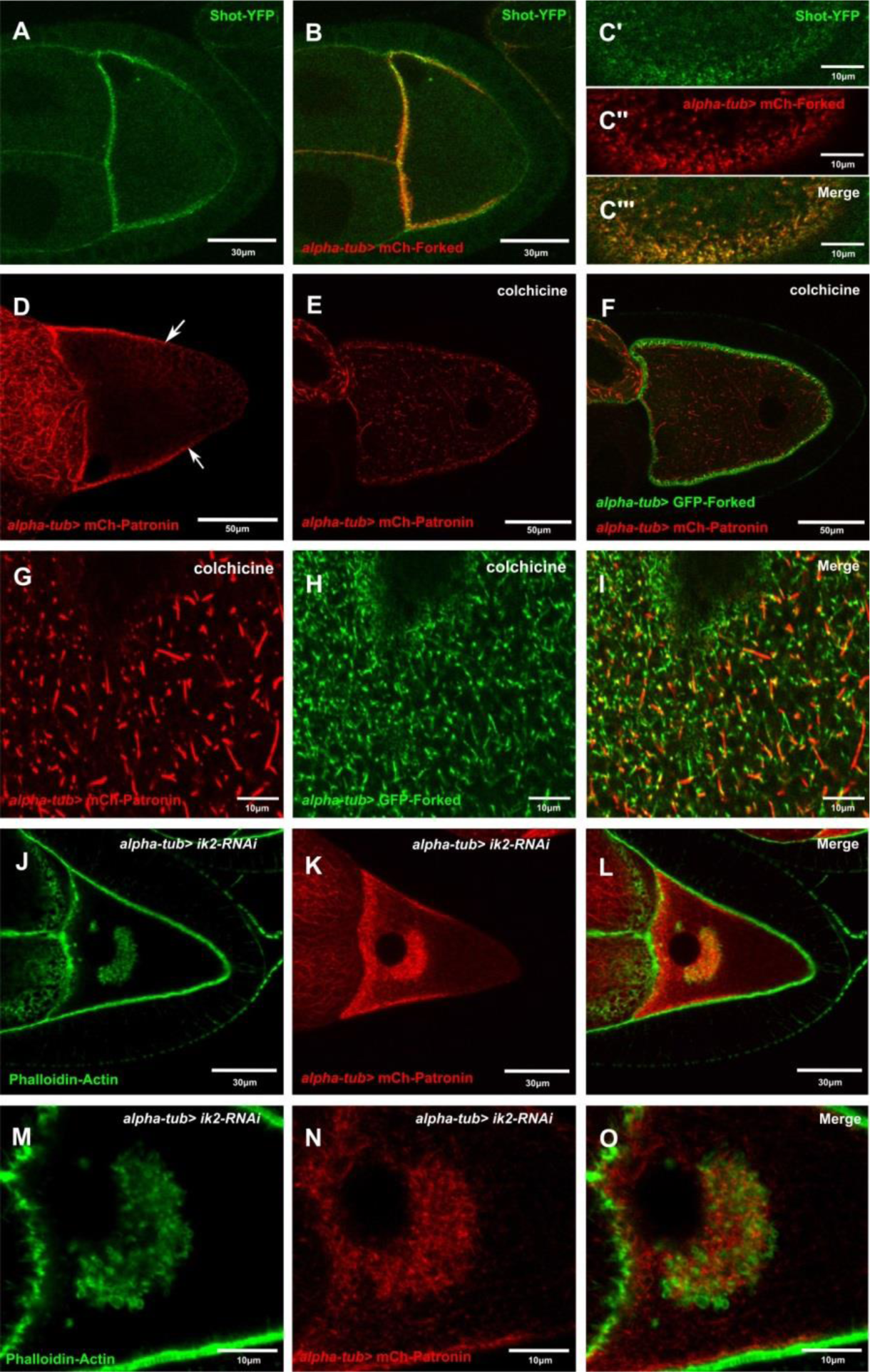
Forked is associated with ncMTOCs. Confocal image of stage 9 egg chamber expressing both (a) Shot-YFP, (b) *alpha-tub>* mCherry-Forked and (C′-C′′′) rectangle slices from the cortex region of egg chamber expressing both (c′) Shot-YFP and (C′′) *alpha-tub>* mCherry-Forked, (C′′′) merged image. (D) Confocal image of stage 10 egg chamber expressing *alpha-tub>* mCherry-Patronin. (E-I) Confocal images of stage 10 egg chambers expressing (F) both *alpha-tub>* mCherry-Patronin (E) and *alpha-tub>* GFP-Forked from the flies fed with colchicine. Confocal images of cortex region from the stage 10 (G-I) egg chambers (D-F). (I) Merged image. (J-L) Confocal images of stage 10 egg chambers from flies expressing both (J) *Ik2 RNAi* and (K) *alpha-tub>* mCherry-Patronin, stained with Phalloidin for actin in green (J). (M-O) Confocal images of stage 10 egg chambers (J-L). (L) and (O) are merged images. Arrows in (D) are pointing towards the limit of the asymmetrical microtubule network marked by mCherry-Patronin.

## Discussion

We previously found that Spn-F, together with IK2, plays a role in both oocyte polarity maintenance (Amsalem et al., 2013; Dubin-Bar et al., 2008) and bristle development (Otani et al., 2015). We, therefore, wanted to test whether other factors important for bristle formation also assume a role in oogenesis. Forked is an actin cross-linker protein that is required for the formation of actin bundles during bristle development (Petersen et al., 1994). We found that *forked* knock-out females were fertile and showed no obvious defects in the localization of either Grk or Stau (Fig. 1B, D). These results demonstrated that the *forked* gene is not required for oocyte development.

Oocyte polarity in *Drosophila* is established in several steps, which involve dynamic changes in the microtubule network. Initially, centrosomes migrate from nurse cells towards the oocyte, accumulating at the posterior end of the oocyte nucleus, where a new MT-organizing center forms. All MTs are nucleated at this posterior MT-organizing center in stage 1 to stage 6 egg chambers (Bolivar et al., 2001; Mahowald and Strassheim, 1970). During the same stages, the asymmetric Forked network that we described here also assembles at the posterior end of the oocyte. In egg chambers at around stage 6/7, a as yet unknown signal from the posterior follicle cells serves to disassemble this posterior MTOC in response to Gurken to EGFR signaling. The formation of new MTs is subsequently initiated at the anterior and lateral cortexes, leading to a reversal of MT polarity in the oocyte. This new polarity, whereby MT minus ends are anchored at the anterior and lateral cortexes, is crucial for the localization of axis determinants (Theurkauf et al., 1992). Significantly, the asymmetric Forked network that we described here also undergoes a posterior to anterior transition. We have shown that the asymmetric distribution of this Forked network depends on MTs, using *gurken* mutants, *par-1* mutants, and colchicine treatment. Indeed, we found that Forked co-localized with both Shot and Patronin foci, and that abnormal ectopic actin clogs are associated with ncMTOCs in the oocyte. Thus, the finding that Forked is associated with ncMTOCs explains the sensitivity of the asymmetric Forked network to conditions that impair the MT network.

The timing and distribution of a polarized microtubule network during mid-oogenesis and the localization of the asymmetrical Forked-marked filamentous network are highly similar. Moreover, the the Forked network is dependent on polarized MTs, as revealed using *par-1* and *grk* mutants. These observations, together with the fact that Forked is associated with ncMTOC, support the use of GFP-Forked as a novel genetic reporter to study *Drosophila* oocyte polarity.

## Materials and Methods

### *Drosophila* stocks

The following mutant and transgenic flies were used (See FlyBase for reference): *f* 36a (Hoover et al., 1993), *grk^2B^ 6* (Neuman-Silberberg and Schupbach, 1993), *grk*^*ΔFRT*^ (Lan et al., 2010), *par-1*^*W3*^ (Shulman et al., 2000), *par-1*^*6323*^ (Shulman et al., 2000), mCherry-Patronin and Shot-YFP (Nashchekin et al., 2016). *shot-RNAi* (#41858), *Df(1)BSC584* (#25418) and *ik2-RNAi* (#35266) were obtained from the Bloomington Stock Center. Germline expression was performed with P{*matα4-* GAL4-VP16} V37 (hereafter referred to as *alpha-tub*), also obtained from the Bloomington Stock Center. Ovaries of the following genotypes were analyzed: (1) *alpha-tub>* GFP-Forked, (2) *alpha-tub>* mCherry-Forked, (3) *grk*^*2B6*^/*grk*^*ΔFRT*^; *alpha-tub>* GFP-Forked, (4) *par-1*^*w3*^/ *par-1^6323^; alpha-tub>* GFP-Forked, (5) *alpha-tub>* mCherry-Forked */shot-RNAi,* (6 *) alpha-tub-2>* mCherry-Forked */Shot-YFP* and (7) *alpha-tub>* mCherry-Patronin; GFP-Forked.

### Transgenic flies

The *forked* gene contains nine alternative splice forms. In this study, we mainly used the *forked-C* isoform, which was studied before (Hoover et al., 1993; Petersen et al., 1994). However, since it was shown that extra copies of *forked* affected bristle morphology (Petersen et al., 1994), we amplified a truncated form of the *forked-C* isoform from pupal cDNA using the following forward primer: 5’- GGAGTTCGTGACCGCCGCCGGGATCACTCTCGGCATGGACGAGCTGTACAAGTCTAGAATGACCACAAGTCTGACCTC-3’ and reverse primer - **5′** TATCAAGCTCCTCGAGTTAACGTTACGTTAACGTTAACGTTCGAGGTCGACTCTAGATCAGAGCAGCTTGGCTTTC-3′.

To make N-terminal GFP- or mCherry-Forked fusions using plasmids pUASp-GFP::Forked and pUASp-mCherry::Forked, DNA for GFP and mCherry was cloned into plasmid pUASp using the *Kpn*I and *Xba*I restriction sites by Gibson assembly (hereafter, the constructs are designated as plasmids pUASp-GFP and pUASp-mCherry). To make an N-terminal GFP-Forked fusion using plasmid pUASt-GFP::Forked, GFP was cloned into plasmid pUASt using the *Kpn*I and *Xba*I restriction sites (hereafter, the construct is designated as plasmid pUASt-GFP). *forked-C* was then cloned into plasmids pUASp-GFP, pUASt-GFP and pUASpmCherry using the *Xba*I restriction sites. P-element-mediated germline transformation of these constructs was carried out by BestGene (Chino hills, C.A).

For generating *forked* knock out flies, appropriate guide RNA sequences were identified (Table S3) at http://tools.flycrispr.molbio.wisc.edu/targetFinder/ and cloned into plasmid pU6-BbsI-chiRNA. Then, 1 Kb sequences stretches upstream and downstream of *forked* was cloned into the donor pHD-DsRed-attP vector. Finally, injection of both vectors and fly screening was carried out by BestGene (Chino hills, C.A).

### Fertility assay

Three virgin females of the respective genotypes were mated with two WT males in a vial containing yeast for two days. Matings were performed in triplicate for each genotype. The flies were transferred to new vials containing fresh yeast for one day to lay eggs. The flies were discarded and the progeny resulting from the eggs after 10 days at 25°C were collected and counted. From each vial, the number of progeny per female, and the average number and standard deviation of progeny per genotype were calculated. Finally, a percentage of relative fertility was calculated (Spracklen et al., 2014).

### Drug Treatment

Flies that had been starved for two hours were fed 200 μg/ml colchicine for 16 h. The ovaries were dissected in PBS and fixed in 4% paraformaldehyde (PFA), followed by Alexa Fluor 568 phalloidin staining for actin.

### Immunohistochemistry

Ovaries were dissected in PBS and fixed for 10 min in 4% PFA. For actin staining, Oregon green 488 and Alexa Fluor 568 phalloidin dyes were used (Life Technologies). For antibody staining, anti-grk (1:50) and anti-staufen (1:500) antibodies were used. All images were taken on an Olympus FV1000 laser-scanning confocal microscope.

## Acknowledgement

We thank Trudi Schüpbach for generating flies stocks and for valuable suggestions. Also, we thank VDRC Austria and the Bloomington Stock Center for generously providing fly strains.

## Competing interests

No competing interest declared.

## Funding

This work was supported by the Israel Science Foundation (ISF) (278/16) (to U.A.).

---

**S1:**
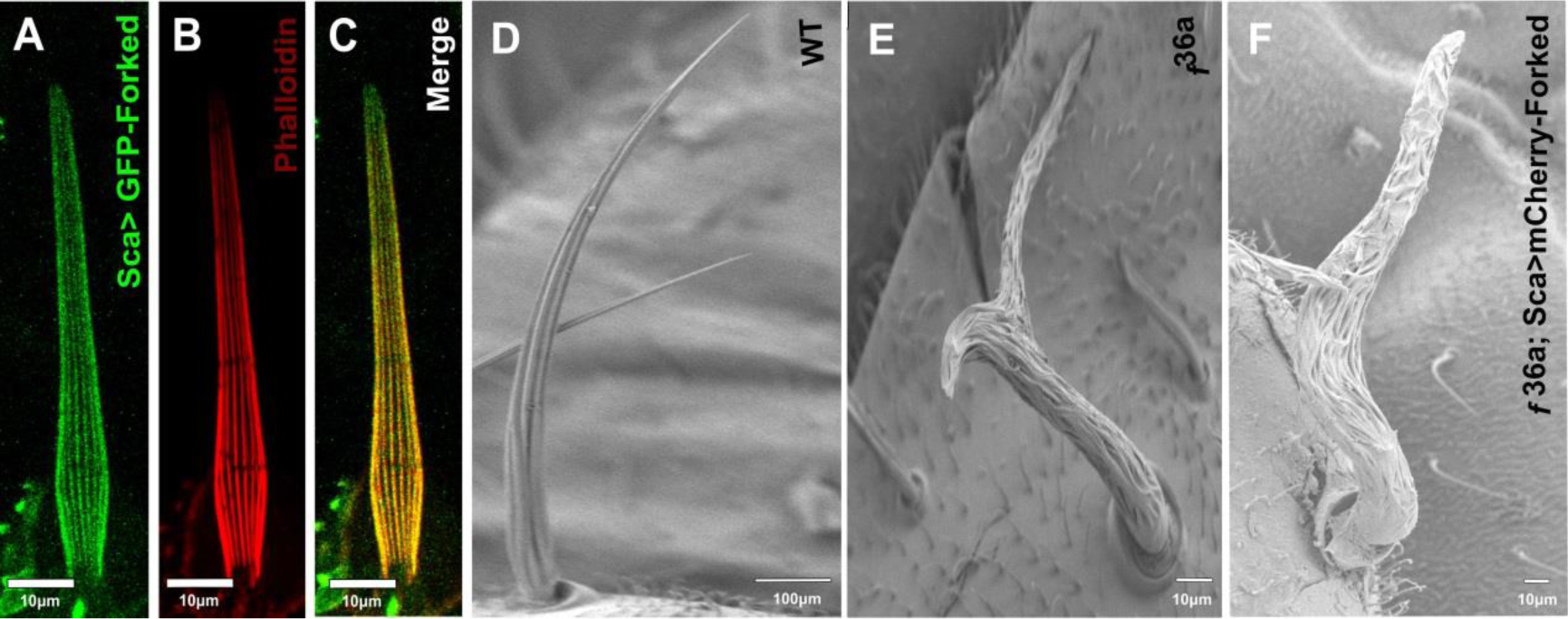
Truncate form of GFP tagged Forked isoform C fails to rescue the *forked* mutant bristle phenotype. Confocal Z-series projections of 40 hrs APF bristle from WT fly expressing (A) *Sca* > GFP-Forked truncate form of isoform C is stained with phalloidin for actin in red (B). (C) Merged image. Scanning electron micrograph of adult bristle from flies of (D) wild type, (E) *f*^*36a*^ showing severe morphological defects and (F) *f*^*36a*^; *Sca* > UAS-mCherry-Forked truncate form of isoform C showing defects similar to that of *f*^*36a*^.

**S2:**
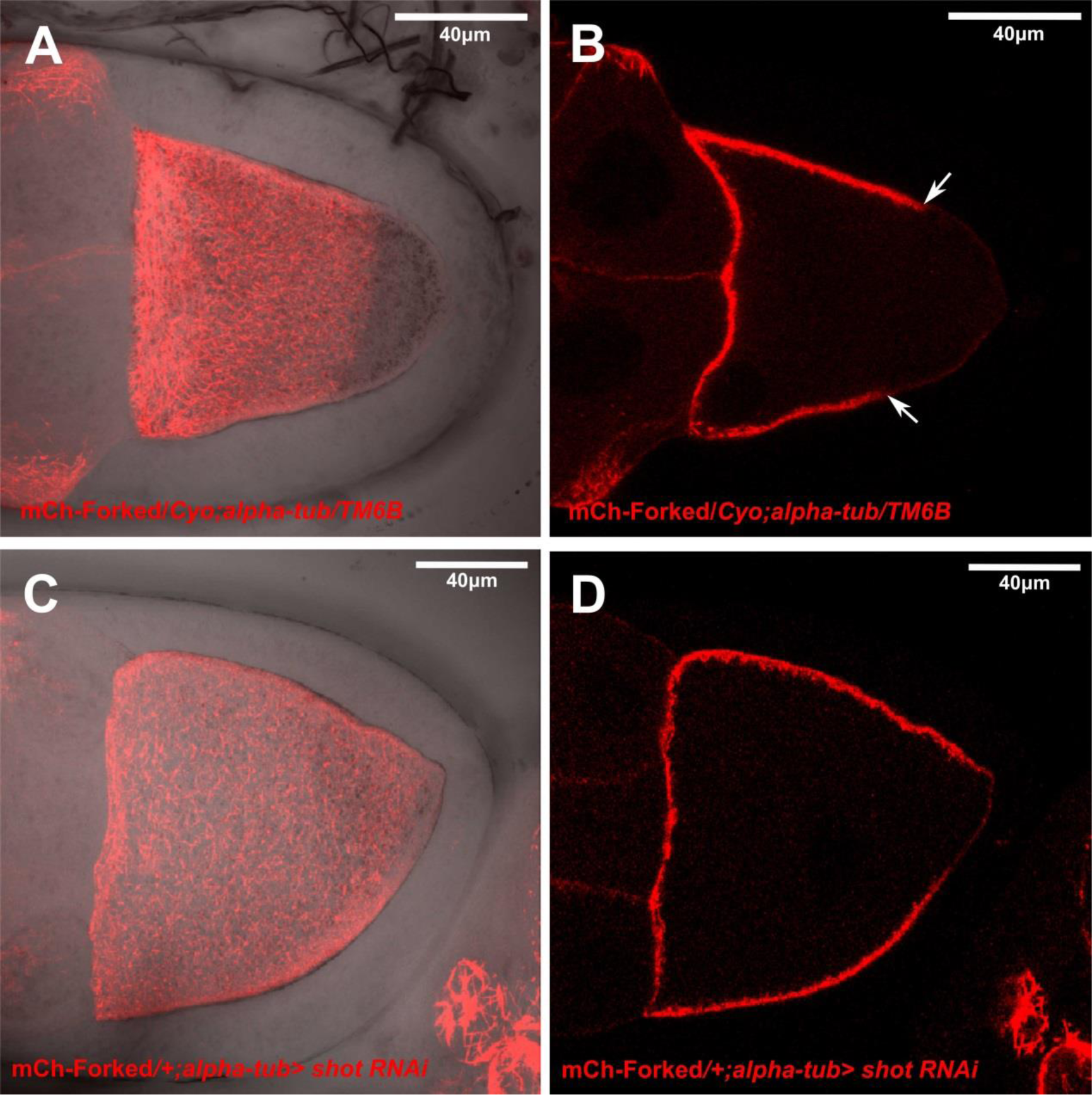
Asymmetric Forked network depends on Shot-Stop. Confocal Z-series projections merged with DIC of stage 10 (A) egg chambers from flies expressing *alpha-tub>* mCherry-Forked and (C) expressing both *Shot RNAi* and *alpha-tub>* mCherry-Forked. (B) and (D) are confocal slice from the image (A) and (C), respectively. Arrows in (B) is pointing towards the limit of the asymmetric network marked by mCherry-Forked.

**Table S1:** forked null allele (CRISPR KO) has no effect on female fertility.

| WT |  |  |  |  |  |
| --- | --- | --- | --- | --- | --- |
| Vials no | Number of progeny from a 24h egg collection | Number of progeny per female | Avg | STD | % relative fertility |
| 1 | 59 | 55 | 55 | 5 | 100 |
| 2 | 65 | 50 |  |  |  |
| 3 | 62 | 60 |  |  |  |
| *forked KO/forked KO* |  |  |  |  |  |
| Vials no | Number of progeny from a 24h egg collection | Number of progeny per female | Avg | STD | % relative fertility |
| 1 | 59 | 46 | 46 | 6 | 83.6 |
| 2 | 62 | 40 |  |  |  |
| 3 | 50 | 52 |  |  |  |

**Table S2:** Overexpression of Forked in the germline has no effect on female fertility.

| *alpha-tub*-GAL4-VP16/*alpha-tub*- GAL4-VP16 |  |  |  |  |  |
| --- | --- | --- | --- | --- | --- |
| Vials no | Number of progeny from a 24h egg collection | Number of progeny per female | Avg | STD | % relative fertility |
| 1 | 71 | 23 | 23 | 3.05505 | 100 |
| 2 | 63 | 21 |  |  |  |
| 3 | 81 | 27 |  |  |  |
| *alpha-tub*> GFP-Forked / *alpha-tub*> GFP-Forked |  |  |  |  |  |
| Vials no | Number of progeny from a 24h egg collection | Number of progeny per female | Avg | STD | % relative fertility |
| 1 | 39 | 13 | 19 | 7.21 | 82.6 |
| 2 | 51 | 17 |  |  |  |
| 3 | 81 | 27 |  |  |  |

**Table S3.**
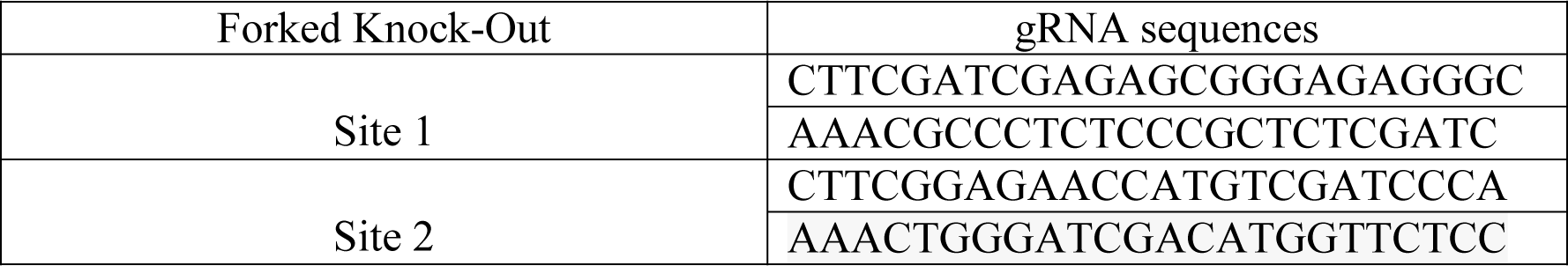
Guide RNA sequences for forked knock out.

## Reference

Abdu, U., Bar, D. and Schupbach, T. (2006). spn-F encodes a novel protein that affects oocyte patterning and bristle morphology in Drosophila. Development 133, 1477–1484.

Amsalem, S., Bakrhat, A., Otani, T., Hayashi, S., Goldstein, B. and Abdu, U. (2013). Drosophila oocyte polarity and cytoskeleton organization require regulation of Ik2 activity by Spn-F and Javelin-like. Mol Cell Biol 33, 4371–4380.

Bitan, A., Guild, G. M., Bar-Dubin, D. and Abdu, U. (2010). Asymmetric microtubule function is an essential requirement for polarized organization of the Drosophila bristle. Mol Cell Biol 30, 496–507.

Bolivar, J., Huynh, J. R., Lopez-Schier, H., Gonzalez, C., St Johnston, D. and Gonzalez-Reyes, A. (2001). Centrosome migration into the Drosophila oocyte is independent of BicD and egl, and of the organisation of the microtubule cytoskeleton. Development 128, 1889–1897.

Cant, K., Knowles, B. A., Mooseker, M. S. and Cooley, L. (1994). Drosophila singed, a fascin homolog, is required for actin bundle formation during oogenesis and bristle extension. J Cell Biol 125, 369–380.

Doerflinger, H., Benton, R., Torres, I. L., Zwart, M. F. and St Johnston, D. (2006). Drosophila anterior-posterior polarity requires actin-dependent PAR-1 recruitment to the oocyte posterior. Curr Biol 16, 1090–1095.

Driever, W. and Nusslein-Volhard, C. (1988a). The bicoid protein determines position in the Drosophila embryo in a concentration-dependent manner. Cell 54, 95–104.

Driever, W. and Nusslein-Volhard, C. (1988b). A gradient of bicoid protein in Drosophila embryos. Cell 54, 83–93.

Dubin-Bar, D., Bitan, A., Bakhrat, A., Amsalem, S. and Abdu, U. (2011). Drosophila javelin-like encodes a novel microtubule-associated protein and is required for mRNA localization during oogenesis. Development 138, 4661–4671.

Dubin-Bar, D., Bitan, A., Bakhrat, A., Kaiden-Hasson, R., Etzion, S., Shaanan, B. and Abdu, U. (2008). The Drosophila IKK-related kinase (Ik2) and Spindle-F proteins are part of a complex that regulates cytoskeleton organization during oogenesis. BMC Cell Biol 9, 51.

Ephrussi, A., Dickinson, L. K. and Lehmann, R. (1991). Oskar organizes the germ plasm and directs localization of the posterior determinant nanos. Cell 66, 37–50.

Gonzalez-Reyes, A., Elliott, H. and St Johnston, D. (1995). Polarization of both major body axes in Drosophila by gurken-torpedo signalling. Nature 375, 654–658.

Hoover, K. K., Chien, A. J. and Corces, V. G. (1993). Effects of transposable elements on the expression of the forked gene of Drosophila melanogaster. Genetics 135, 507–526.

Huynh, J. R. and St Johnston, D. (2004). The origin of asymmetry: early polarisation of the Drosophila germline cyst and oocyte. Curr Biol 14, R438–49.

Lan, L., Lin, S., Zhang, S. and Cohen, R. S. (2010). Evidence for a transport-trap mode of Drosophila melanogaster gurken mRNA localization. PLoS One 5, e15448.

Legent, K., Tissot, N. and Guichet, A. (2015). Visualizing Microtubule Networks During Drosophila Oogenesis Using Fixed and Live Imaging. Methods Mol Biol 1328, 99–112.

Lundh, T., Udagawa, J., Hanel, S. E. and Otani, H. (2011). Cross- and triple-ratios of human body parts during development. Anat Rec (Hoboken) 294, 1360–1369.

Mahowald, A. P. and Strassheim, J. M. (1970). Intercellular migration of centrioles in the germarium of Drosophila melanogaster. An electron microscopic study. J Cell Biol 45, 306–320.

Nashchekin, D., Fernandes, A. R. and St Johnston, D. (2016). Patronin/Shot Cortical Foci Assemble the Noncentrosomal Microtubule Array that Specifies the Drosophila Anterior-Posterior Axis. Dev Cell 38, 61–72.

Neuman-Silberberg, F. S. and Schupbach, T. (1993). The Drosophila dorsoventral patterning gene gurken produces a dorsally localized RNA and encodes a TGF alpha-like protein. Cell 75, 165–174.

Neuman-Silberberg, F. S. and Schupbach, T. (1996). The Drosophila TGF-alpha-like protein Gurken: expression and cellular localization during Drosophila oogenesis. Mech Dev 59, 105–113.

Otani, T., Oshima, K., Kimpara, A., Takeda, M., Abdu, U. and Hayashi, S. (2015). A transport and retention mechanism for the sustained distal localization of Spn-FIKKepsilon during Drosophila bristle elongation. Development 142, 3612.

Parton, R. M., Hamilton, R. S., Ball, G., Yang, L., Cullen, C. F., Lu, W., Ohkura, H. and Davis, I. (2011). A PAR-1-dependent orientation gradient of dynamic microtubules directs posterior cargo transport in the Drosophila oocyte. J Cell Biol 194, 121–135.

Petersen, N. S., Lankenau, D. H., Mitchell, H. K., Young, P. and Corces, V. G. (1994). forked proteins are components of fiber bundles present in developing bristles of Drosophila melanogaster. Genetics 136, 173–182.

Roth, S., Neuman-Silberberg, F. S., Barcelo, G. and Schupbach, T. (1995). cornichon and the EGF receptor signaling process are necessary for both anterior-posterior and dorsal-ventral pattern formation in Drosophila. Cell 81, 967–978.

Shapiro, R. S. and Anderson, K. V. (2006). Drosophila Ik2, a member of the I kappa B kinase family, is required for mRNA localization during oogenesis. Development 133, 1467–1475.

Shulman, J. M., Benton, R. and St Johnston, D. (2000). The Drosophila homolog of C. elegans PAR-1 organizes the oocyte cytoskeleton and directs oskar mRNA localization to the posterior pole. Cell 101, 377–388.

Spracklen, A. J., Fagan, T. N., Lovander, K. E. and Tootle, T. L. (2014). The pros and cons of common actin labeling tools for visualizing actin dynamics during Drosophila oogenesis. Dev Biol 393, 209–226.

St Johnston, D., Beuchle, D. and Nusslein-Volhard, C. (1991). Staufen, a gene required to localize maternal RNAs in the Drosophila egg. Cell 66, 51–63.

Theurkauf, W. E. (1994). Microtubules and cytoplasm organization during Drosophila oogenesis. Dev Biol 165, 352–360.

Theurkauf, W. E., Smiley, S., Wong, M. L. and Alberts, B. M. (1992). Reorganization of the cytoskeleton during Drosophila oogenesis: implications for axis specification and intercellular transport. Development 115, 923–936.

